# Design of linear and cyclic peptide binders of different lengths from protein sequence information

**DOI:** 10.1101/2024.06.20.599739

**Authors:** Qiuzhen Li, Efstathios Nikolaos Vlachos, Patrick Bryant

**Affiliations:** The Department of Molecular Biosciences, The Wenner-Gren Institute, Stockholm University, Svante Arrhenius väg 20C, 114 18 Stockholm, Sweden; Science for Life Laboratory, 172 21, Solna, Sweden

## Abstract

Structure prediction technology has revolutionised the field of protein design, but key questions such as how to design new functions remain. Many proteins exert their functions through interactions with other proteins, and a significant challenge is designing these interactions effectively. While most efforts have focused on larger, more stable proteins, shorter peptides offer advantages such as lower manufacturing costs, reduced steric hindrance, and the ability to traverse cell membranes when cyclized. However, less structural data is available for peptides and their flexibility makes them harder to design. Here, we present a method to design both novel linear and cyclic peptide binders of varying lengths based solely on a protein target sequence. Our approach does not specify a binding site or the length of the binder, making the procedure completely blind. We demonstrate that linear and cyclic peptide binders of different lengths can be designed with nM affinity in a single shot, and adversarial designs can be avoided through orthogonal *in silico* evaluation, tripling the success rate. Our protocol, *EvoBind2* is freely available https://github.com/patrickbryant1/EvoBind.

## Introduction

Protein design is in its golden age due to the advent of accurate structure prediction technology [1–3]. While single protein structures can now be designed with high accuracy [4,5], designing new functions, such as binding, without relying on structural information remains challenging [6,7]. Additionally, there are numerous issues to address before designed proteins can benefit society, including scalable production. However, shorter proteins, such as peptides, can bind with high affinity to a diverse set of targets [8,9] and are cost-effective to produce via conventional chemical synthesis.

The success rate in predicting the structure of protein complexes using AlphaFold2 (AF) and AlphaFold-Multimer (AFM) is approximately 60% [2,3]. However, for protein-peptide complexes, AF reports a high false negative rate (84/96) [10,11]. While AF can identify true binders with high confidence, it misses most binders [12]. Therefore, the challenge in peptide binder design is to find a sequence likely to bind that AF can also accurately predict. The subset of such sequences among the 20^L^ possibilities may be very small (or even absent), necessitating a highly efficient and robust search process.

Currently, most protein design workflows consist of two separate steps. Methods like RFdiffusion [5] and backpropagation through AF [13] have been used to design backbone scaffolds for both single and multi-chain proteins. The sequences for these scaffolds are then redesigned using ProteinMPNN [14]. Alternatively, a joint search over a relaxed sequence-structure space [15,16] can be performed, allowing for simultaneous design of sequence and structure. This approach may increase the likelihood of success as the joint sequence-structure space encompasses all possible sequences and structures.

AF relies on coevolution to produce accurate predictions and has been trained to be robust to minor changes in a multiple sequence alignment (MSA), thereby learning the redundancy between sequence and structure [17]. For design tasks, where no MSA or coevolution exists and only a single sequence is used [14,18], AF must infer the structure from the single sequence. Small changes such as single mutations can now have a significant impact on the outcome and it has been shown that AF can distinguish such changes in the binding residues of peptides [10,16].

Based on these principles, we developed a method to generate peptide binders for a target protein using only the protein target sequence. We allow AF to select a suitable binding site, binding sequence, and structure without providing any predefined information. To avoid adversarial designs and increase the success rate, we incorporate a modified version of AFM [2] for evaluation. Additionally, we implement a cyclic offset [19], [20], demonstrating that cyclic peptide binders can be designed to match the efficacy of linear designs using the same input: a single protein target sequence.

## Results

### Linear and Cyclic peptide binder design

Many proteins exert their functions through interactions. The ability to target specific proteins through binding is essential for numerous biotechnological applications such as fluorescent labelling [21], payload delivery [22], targeted degradation [23] and receptor activation/inhibition [24,25]. Current methods for protein binder design rely on knowledge of the target protein structure and binding site [14] and even that of the binder [7] **(Figure 1a).** For novel targets, information about (**1)** target structure, (**2)** target site (residues), or (**3)** binder size is often unknown. In addition, most methods design large binders with well defined structure, but shorter peptides are less sterically hindered and are cost-effective to produce via conventional chemical synthesis. Therefore, we set out to design novel peptide binders for a target protein using only the target protein sequence **(Figure 1b)**.

**Figure 1.**
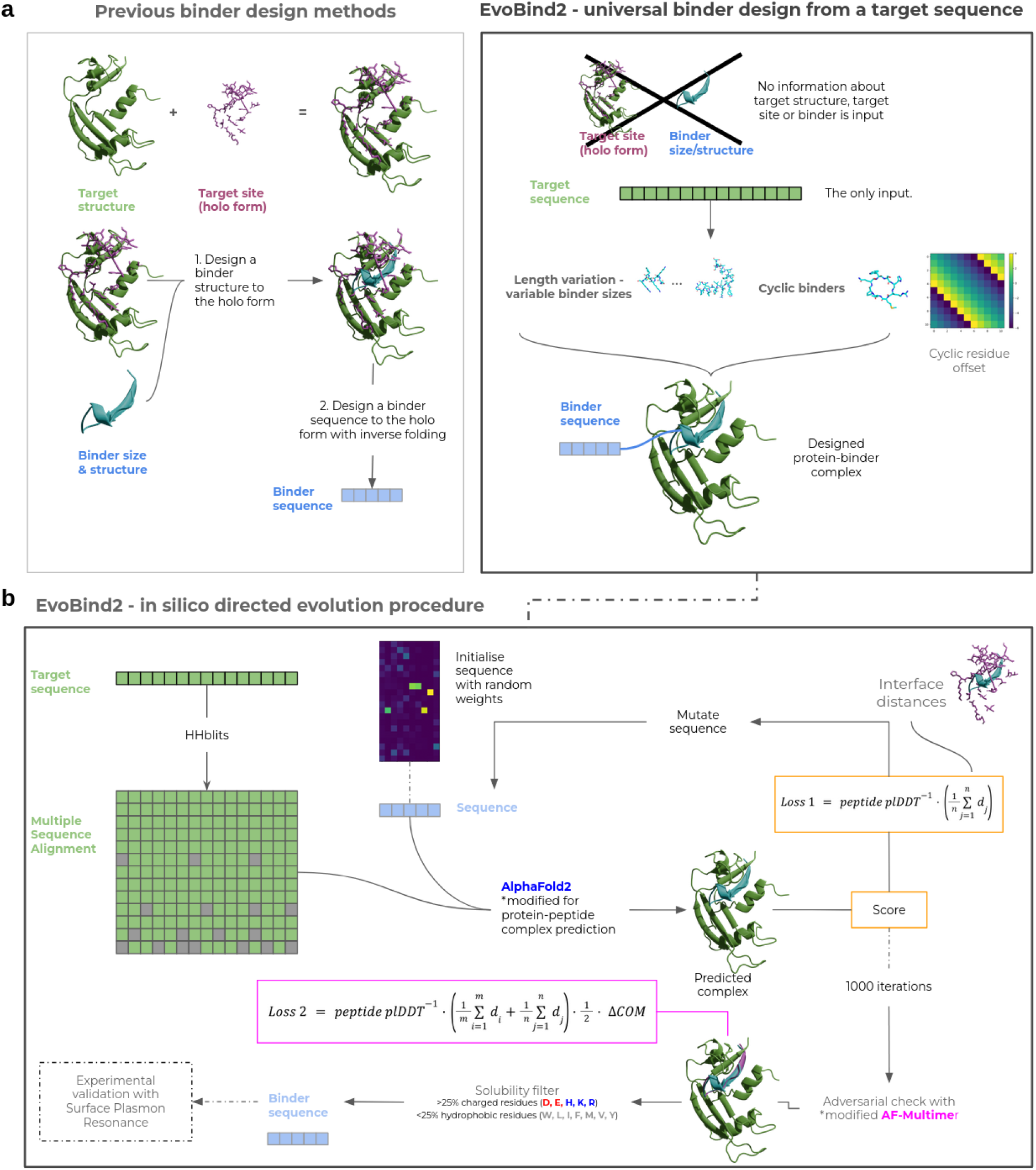
**a)** Other methods for binder design rely on structural knowledge of target structures, binding sites in holo form and binders. This hampers the development of novel binders as binding information is required as input. In contrast, EvoBind2 is based solely on the sequence of the protein target. This makes EvoBind2 fully flexible and a binding sequence and protein-peptide complex structure is generated and evaluated simultaneously. The user can define the length (number of amino acids) of the binder and we find that Evobind can adapt the design to various peptide lengths (Figure 2). By implementing a cyclic offset, we also enable cyclic peptide binder design offering potential cell membrane traversal and increased peptide half-lives. **b)** Procedure for the design of peptide binders with EvoBind2. The target protein sequence is the only input to the design pipeline. No information about the binding site or peptide length is provided. The target sequence is searched with HHblits against Uniclust30, resulting in a multiple sequence alignment (MSA).

**Figure 2.**
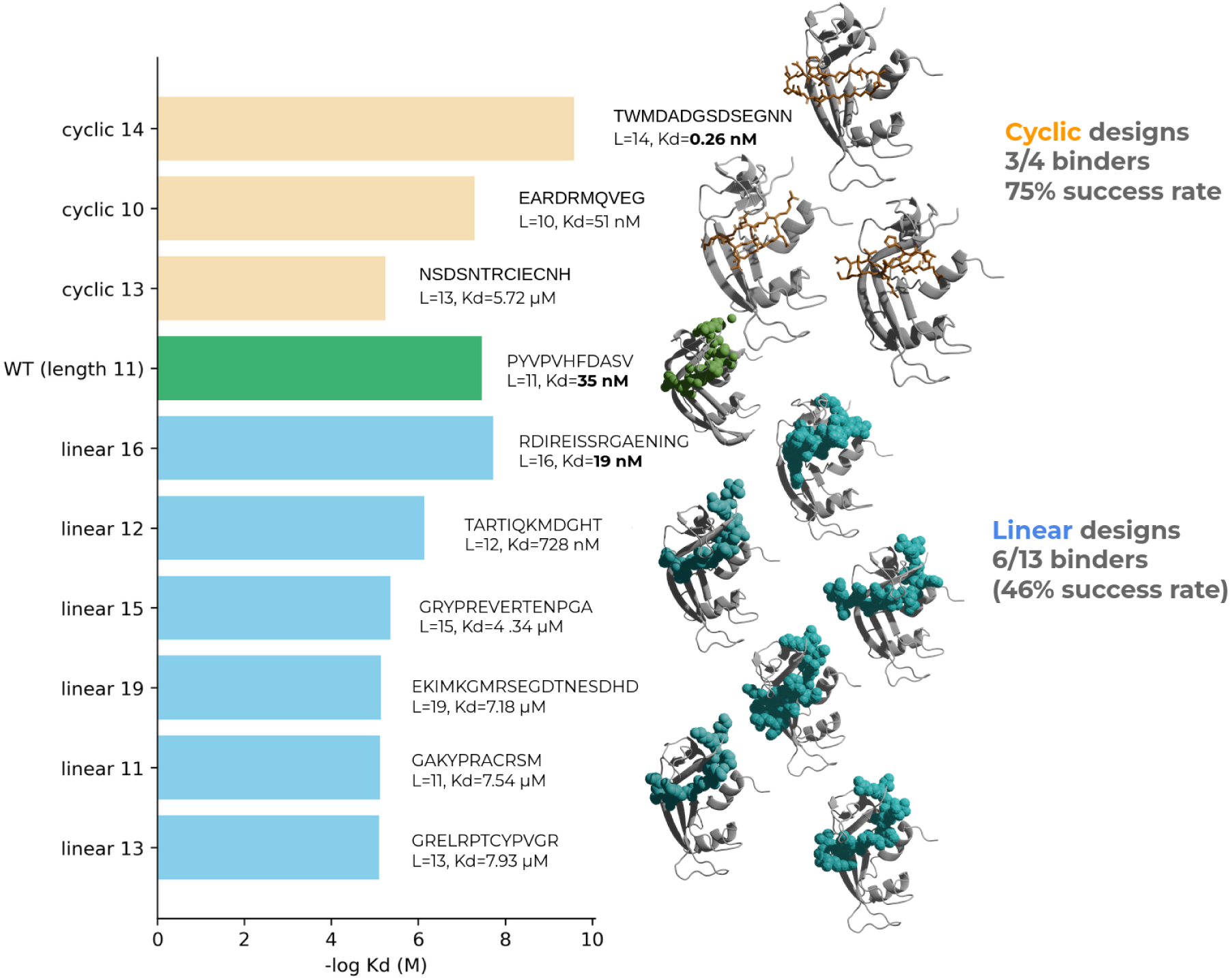
**a)** Affinity (Kd) and predicted structure for the selected successful designs measured by surface plasmon resonance (SPR, methods). The predicted structures are shown with the target protein in grey and the peptide binder in orange stick format (cyclic) and cyan spheres (linear). The wildtype (WT) linear peptide is shown as green spheres in its native structure from the PDB ID 1SSC: https://www.rcsb.org/structure/1ssc. In the cyclic case, 75% of the evaluated binders have at least μM affinity and the strongest binder has a Kd 137 times lower than the wild type (0.26 vs 35 nM). In the linear case, 6/13 lengths are successful and the best binder has a Kd almost twice that of the wild type (19 vs 35 nM).

To design binders, we create a framework based on protein structure prediction [1], **EvoBind2,** for *in silico* directed evolution [15] (**Figure 1c**). EvoBind2 iteratively mutates a peptide binder sequence to minimise the distance from the predicted positions of the peptide atoms towards any receptor protein atom (interface distances) and maximise the predicted confidence (peptide plDDT [1], Loss 1, equation 1). This approach allows complete freedom in generating a sequence-structure combination for a peptide binder where EvoBind2 deems appropriate. This lack of constraint removes any procedural bias, as preconceived notions about what constitutes a good binding structure, sequence, or site may be suboptimal.

The design pipeline uses only the target protein sequence as input, with no information about the binding site or peptide length. The target sequence is searched against Uniclust30 with HHblits, generating a multiple sequence alignment (MSA). The peptide sequence is randomly initialised, and the resulting sequence is fed into a modified AlphaFold2 (AF) to predict the protein-peptide complex structure. A score (Loss 1) is calculated from the result. If the loss is lower than previously, the new peptide sequence is used as a starting point for introducing a new mutation and the procedure is then repeated. The sequence is subject to 1000 rounds of mutation and the AF predictions are then compared to predictions from a modified AlphaFold-multimer (AFM, Loss 2). This adversarial check is followed by a solubility filter and the final sequences are experimentally evaluated using surface plasmon resonance.

The peptide sequence is randomly initialised with weights in the range 0-1 from a Gumbel distribution and the argmax at each position (Lx20) is used as the initial amino acid sequence. The MSA and the initial peptide sequence is input to a modified version of AlphaFold2 to predict the structure of the protein-peptide complex. A score (Loss 1) is calculated from the result. If the loss is lower than previously, the peptide sequence is used as a starting point for introducing a mutation and the complex prediction is then repeated. The sequence is subject to 1000 rounds of mutation and the AlphaFold2 predictions are then compared to predictions from a modified AlphaFold-multimer (Loss 2). This adversarial check is followed by a solubility filter and the final sequences are experimentally evaluated using surface plasmon resonance.

Using EvoBind2, we designed both linear and cyclic peptide binders of 8-20 residues targeting the semi-synthetic Ribonuclease (PDB ID 1SSC: https://www.rcsb.org/structure/1ssc). We expressed and purified the protein, synthesised the peptides and measured the binding affinity of the designed peptides using surface plasmon resonance (SPR) (Methods). We selected the top sequence for each length (n=13 lengths) in the linear case and the top 5 lengths in the cyclic one, although one of these could not be easily synthesised (Methods). We found that 6 out of 13 (46%) of the designed linear peptide binders displayed μM affinity, with the weakest binder having a dissociation constant (Kd) of 7.93 μM and the strongest a Kd of 19 nM (**Figure 1)**.

For the cyclic designs, we evaluated the top four designs experimentally and lengths 10, 13 and 14 obtained Kds of 51 nM, 5.72 μM and 0.26 nM, respectively (75% success rate, **Figure 1**). Compared to the positive control (Kd = 35 nM, Methods), the strongest linear binder exhibited almost twice and the strongest cyclic 137 times the affinity from a single selected sequence. Additionally, the peptides display a diverse set of sequences and predicted binding modalities, highlighting EvoBind2’s capability to generate varied solutions. We note that neither AF or AFM have seen any structures similar to the designed protein-peptide complexes (especially in the cyclic case), suggesting the capability of EvoBind to generalise to unseen targets.

The loss function used for the optimisation is defined as:

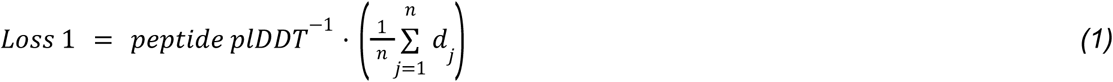

Where the peptide plDDT is the average plDDT over the peptide and dj is the shortest distance between all atoms n in the peptide and any atom in the target protein. The plDDT, the predicted local distance difference test (pLDDT) [26], evaluates the local confidence of predicted protein structures. Higher pLDDT scores indicate higher confidence.

### Adversarial designs

When designing binders to proteins with structure prediction technology (AF), it is possible to obtain designs that exhibit high *in silico* confidence (low loss, equation 1) without being true binders. These sequences that “trick” EvoBind2 are deemed adversarial [27]. To avoid adversarial designs, we modified AlphaFold-multimer [2] (AFM) to predict protein-ligand complexes using only MSA information for the receptor protein and a single sequence for the peptide. This additional validation increases the likelihood of success, as it is unlikely that two different structure prediction networks, trained on different data, would agree while both are incorrect.

To analyse the usefulness of this additional adversarial check, we selected sequences from the linear design where EvoBind2 and AFM disagree - specifically, where the AFM loss (equation 2) is high but the EvoBind2 loss (equation 1) is low (**Figure 3**). We selected the lowest EvoBind2 loss for each length when the AFM loss was >1, obtaining EvoBind2 losses in the range of 0.04-0.1. We synthesised the peptides and evaluated their binding affinity against the target protein using surface plasmon resonance (SPR).

**Figure 3.**
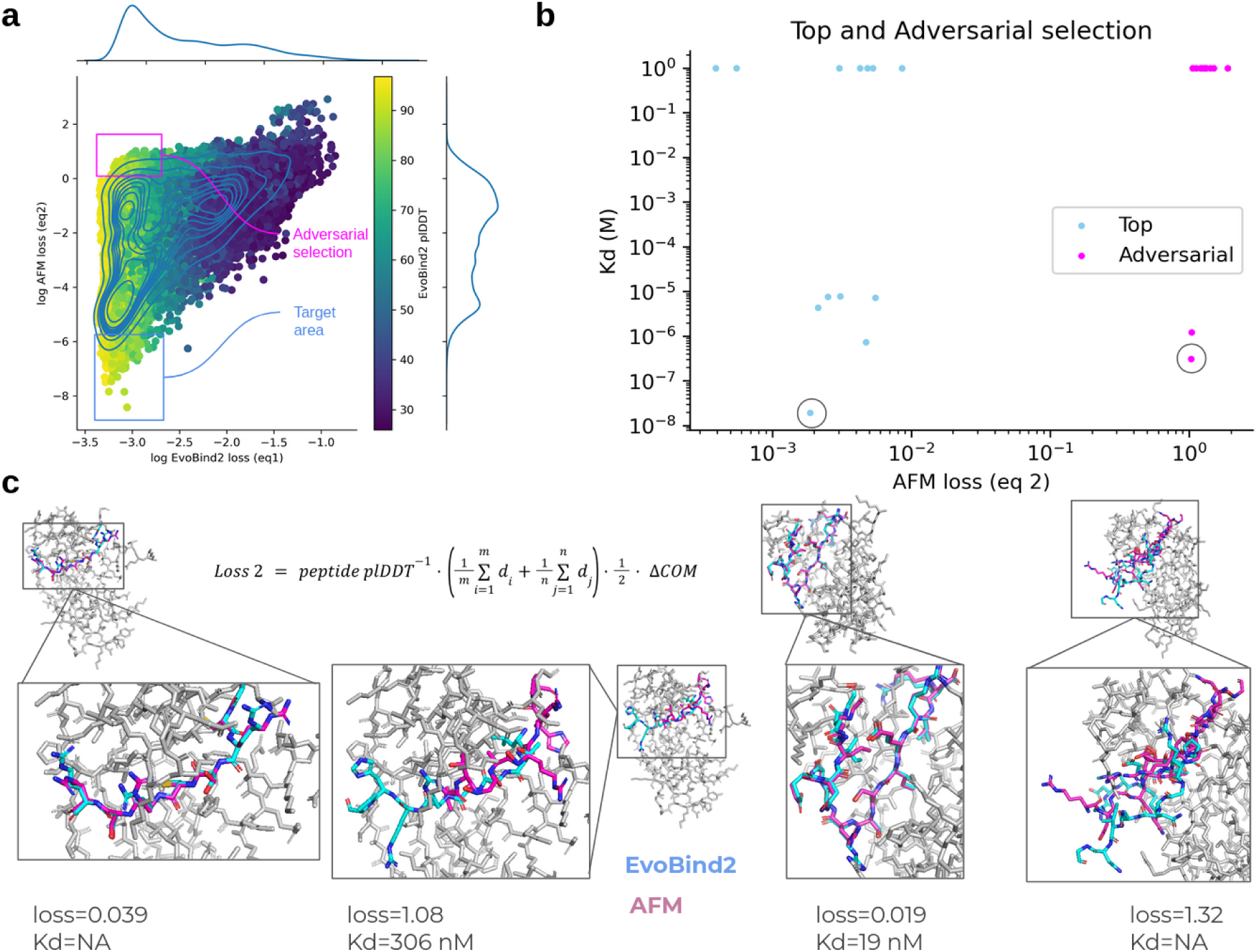
**a)** EvoBind2 loss (equation 1) vs AlphaFold-multimer (AFM) loss (equation 2) on log scale for all linear design runs (n=64935). The points are coloured by the plDDT from EvoBind2 and the density is represented with blue lines. The target area represents cases where both losses are low, and the adversarial selection cases where the EvoBind2 loss is low but the AFM loss is high. The discrepancy between the losses in the adversarial area signifies that those sequences may trick EvoBind2. **b)** Affinity (Kd) vs AFM loss (eq2) for both the top selection (in the target area, Figure 3a) and the adversarial selection (AFM loss>1) for the linear design (n=13). The peptides with no detectable affinity are set to 1 M. The peptides with low AFM loss have lower Kd overall (6/13 with μM affinity) compared to sequences with AFM loss>1 (2/13 with μM affinity). The circled points mark examples shown in c. **c)** Examples of adversarial selections. The predicted structure is shown in stick format with the receptor structures in grey and the predicted peptide structures in cyan/magenta for EvoBind2/AlphaFold-multimer (AFM), respectively. Left; an example adversarial to AFM (length=8) where the Kd is 306 nM although the AFM loss is high and undetectable at a low AFM loss. Right; an example where the adversarial prediction of AFM is accurate (length=16) where the Kd is 19 nM with a low AFM loss and undetectable at a high AFM loss.

We found that only two of the sequences in the adversarial selection displayed detectable affinity compared to six when both losses (equations 1 and 2) were low. This suggests that adversarial selection helps distinguish true binders from false ones, increasing the success rate threefold. Conversely, some sequences seem adversarial to AFM as 2/13 sequences did display affinity (Figure 3c).

The loss for the adversarial selection is defined as:

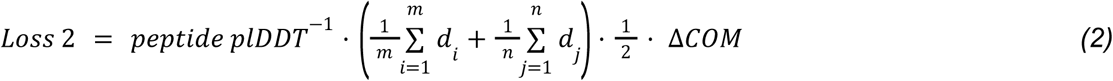

Where the peptide plDDT is the average plDDT over the peptide, di is the shortest distance between all atoms in the receptor target atoms (Cβs within 8 Å between the peptide and the receptor protein as predicted by EvoBind2) m and any atom in the peptide, dj is the shortest distance between all atoms n in the peptide and any atom in the receptor target atoms and ΔCOM is the CA centre of mass distance between the predicted peptides from the design and validation procedures, respectively.

### Affinity and *in silico* metrics

The loss functions (equations 1 and 2) used in this study produce high-affinity binders, but not all selected binders display measurable affinity. To determine whether we can distinguish between true and false binders or predict affinity, we analysed various *in silico* metrics (**Figure 4**). One of the most important metrics used in the loss functions is the predicted local distance difference test (pLDDT) [26], which evaluates the per-residue confidence of predicted protein structures. Higher pLDDT scores indicate higher confidence, but we observe no meaningful correlation between the dissociation constant (Kd) and EvoBind2 (AF) or AFM pLDDT scores (**Figure 4b, 3f**). We also analysed the Predicted Aligned Error (PAE) as a complementary *in silico* metric analogous to the pLDDT. Consistent with the pLDDT findings, we observed no meaningful correlation between Kd and PAE values (**Supplementary Figure 8**).

**Figure 4.**
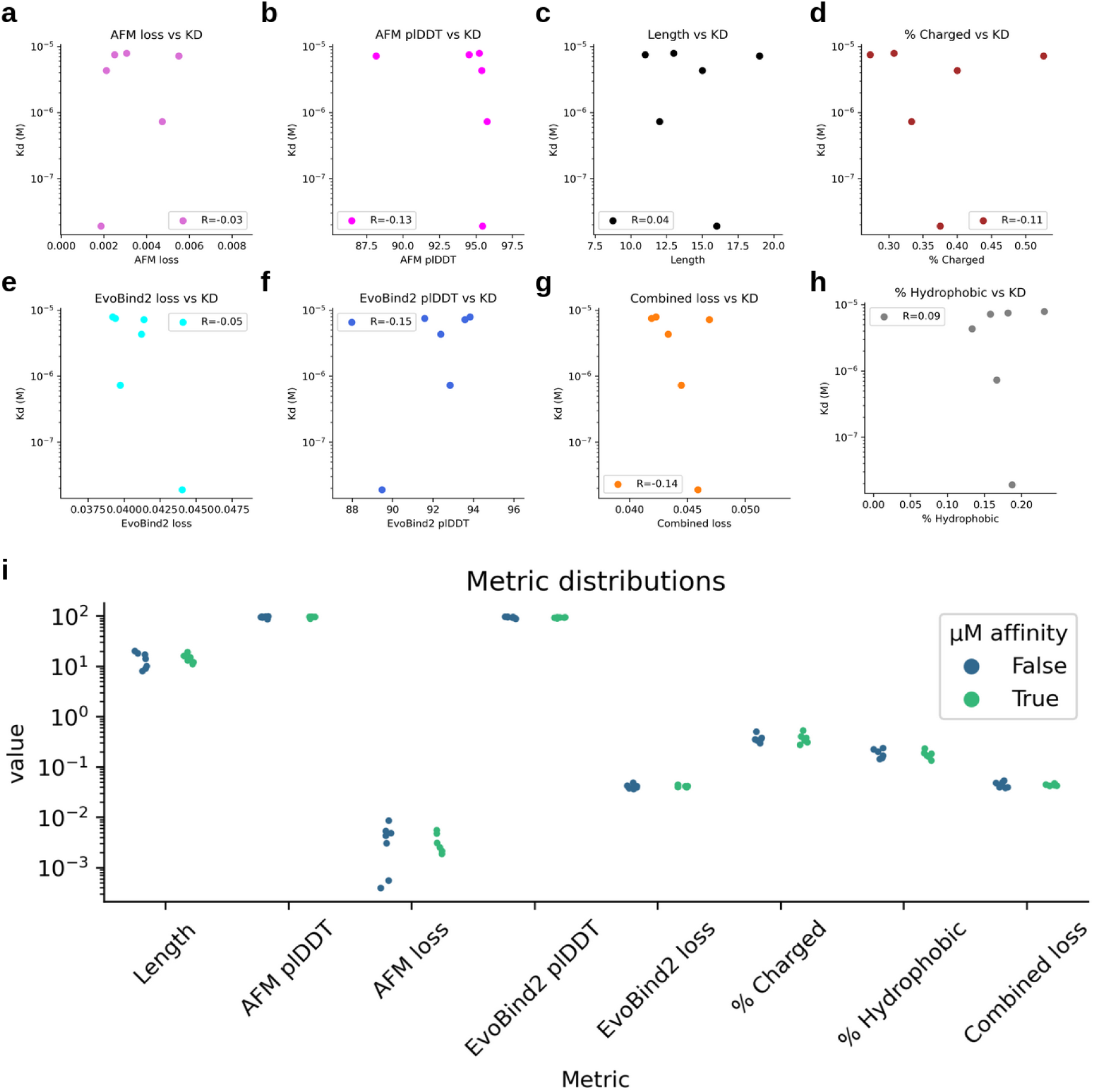
**a-h** Scatter plots showing the affinities of linear peptides (KD) vs *in silico* metrics for successful designs from the top selection (n=6). The Spearman correlation coefficient is annotated in each subfigure. None of the metrics display a meaningful correlation with the measured Kd. **a)** AlphaFold-multimer loss (equation 2) **b)** plDDT from AlphaFold-multimer **c)** the length of designed peptide **d)** the fraction of charged amino acids in the sequence **e)** plDDT from EvoBind2 **f)** the loss from EvoBind2 (equation 1) **g)** the combined loss from EvoBind2 and AlphaFold-multimer (equations 1 and 2) **h)** the fraction of hydrophobic amino acids in the sequence. **i)** Strip plot of the categories in a-h divided by if μM affinity could be measured/not (n=13). No *in silico* metric separates true from false binders.

In addition to generating accurate binders, the peptides must be functional under biological conditions to be useful in biotechnological applications. To ensure this, we applied a simple solubility filter requiring >25% of peptide residues to be charged and <25% to be hydrophobic (Methods). These metrics, however, do not correlate with success in the binder design process (Figure 7c, d) and cannot be used to select for affinity.

The length of a peptide can influence its function, as longer peptides offer greater complexity and potential for diverse interactions. However, we find no relationship between peptide length and Kd (**Figure 4c**), suggesting that the binding residues themselves are what matter most. A comparison of all metrics with the presence or absence of measured μM affinity is displayed in **Figure 4i**, highlighting that *in silico* metrics fail to effectively distinguish between true and false binders. This underlines the complexity of predicting protein-peptide interactions.

## Discussion

AI-based peptide binder design offers the potential to rapidly expand the repertoire of proteins that can be targeted for biotechnological applications. The EvoBind2 framework developed here designs peptide binders to a protein target using only its amino acid sequence. This approach does not require knowledge of binding sites, template structures, or binder sizes making it applicable to novel targets. We observed binding affinities in the range of Kd 5.7 μM to 0.26 nM for cyclic and 7.9 μM to 19 nM for linear peptides, achieving success rates of 75% and 46%, respectively.

Unlike protein structure prediction, which relies on coevolutionary information from multiple sequence alignments (MSAs), *de novo* peptide design lacks this evolutionary context. The absence of coevolutionary information presents a significant challenge when designing new peptides, as the structure must be predicted without these constraints. The finding that successful peptide design is still possible [15] suggests that important protein sequence-structure relationships have been learned independently of coevolution.

One of the key challenges in designing peptide binders is avoiding sequences that have high *in silico* confidence (high pLDDT) but do not bind to the target protein, known as adversarial designs. To address this issue, we adapted AlphaFold-multimer (AFM) [2] to predict protein-ligand complexes using MSA information for the receptor protein and a single sequence for the peptide. This additional validation step, which filters out sequences that appear promising according to EvoBind2 but do not function as true binders, enhances the likelihood of successful linear binder design threefold. The rationale is that it is unlikely for two distinct neural networks, trained on different data, to agree on an incorrect prediction. Previous design methods have not included adversarial checks, which may explain lower success rates observed with other strategies [6,7,16].

The protein targeted in this study, a semi-synthetic Ribonuclease (PDB ID 1SSC: https://www.rcsb.org/structure/1ssc) has a known peptide binder of length 11. To ensure the expressed protein is functional for affinity measurements, we also synthesised this sequence and measured its affinity as a positive control with a resulting Kd of 35 nM. Interestingly, both linear and cyclic binders designed here outperform this control two and 137-fold, respectively, suggesting that EvoBind2 can also be used to improve affinities for targets with known binders.

The predicted structures of the designed peptides show diverse binding modalities (**Figure 2**). However, it is not clear what specific factors contribute to one peptide binding with higher affinity than another (Figure 4). While sequences with a high AFM loss (equation 2) have a lower chance of binding, for sequences with low AFM loss, it remains impossible to distinguish between nM and μM affinity based on predicted confidence metrics alone. The findings highlight the importance of multi-modal validation and emphasises the need for wet-lab validation and careful interpretation when models disagree.

Cyclic peptides offer advantages over linear peptides in terms of stability and potential membrane traversal [28] and several drugs on the market today are cyclic peptides [29]. Interestingly, our results indicate that cyclic peptides can be designed to be comparable or better to linear ones using the same input information. In conclusion, our study demonstrates the potential of EvoBind2 for designing novel peptide binders with high affinity using only a target protein sequence. The lack of a requirement for defined binding sites or binder sizes is crucial for targeting novel proteins and suggests a potential for a rapid increase in the number of proteins that can be targeted for various biotechnological applications.

## Availability

Data: https://zenodo.org/records/13913345

Code: https://github.com/patrickbryant1/EvoBind

## Contributions

PB designed the studies. PB developed and QL executed the binder design procedure. QL performed all laboratory analyses and analysed the results in consultation with PB and ENV. PB wrote the first draft of the manuscript which was later improved by all authors. PB obtained funding.

## Acknowledgements

This study was supported by the SciLifeLab & Wallenberg Data Driven Life Science Program (grant: KAW 2020.0239, P.B). Computational resources were enabled by the supercomputing resource Berzelius provided by National Supercomputer Centre at Linköping University and the Knut and Alice Wallenberg foundation with project ids Berzelius-2023-267, Berzelius-2024-78 and Berzelius-2024-292 (P.B).

The protein purification was facilitated by the Protein Science Facility at Karolinska Institutet, Stockholm, and we would like to thank Dr Henry Ampah-Korsah, Dr Henrik Spåhr, and Dr Tomas Nyman for their assistance.

## Methods

### Design protocol

#### EvoBind2

For designing peptide binders, we use a modified version of EvoBind [15] (**Figure 5**) that does not require a defined target area. This is the first method developed that can generate both sequence and structure from a single protein target sequence (to our knowledge). We design over a relaxed sequence-structure space that is searched by mutating one residue randomly at a time to minimise the loss function:

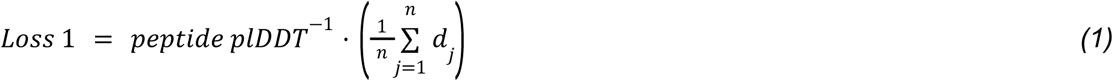

**Figure 5.**
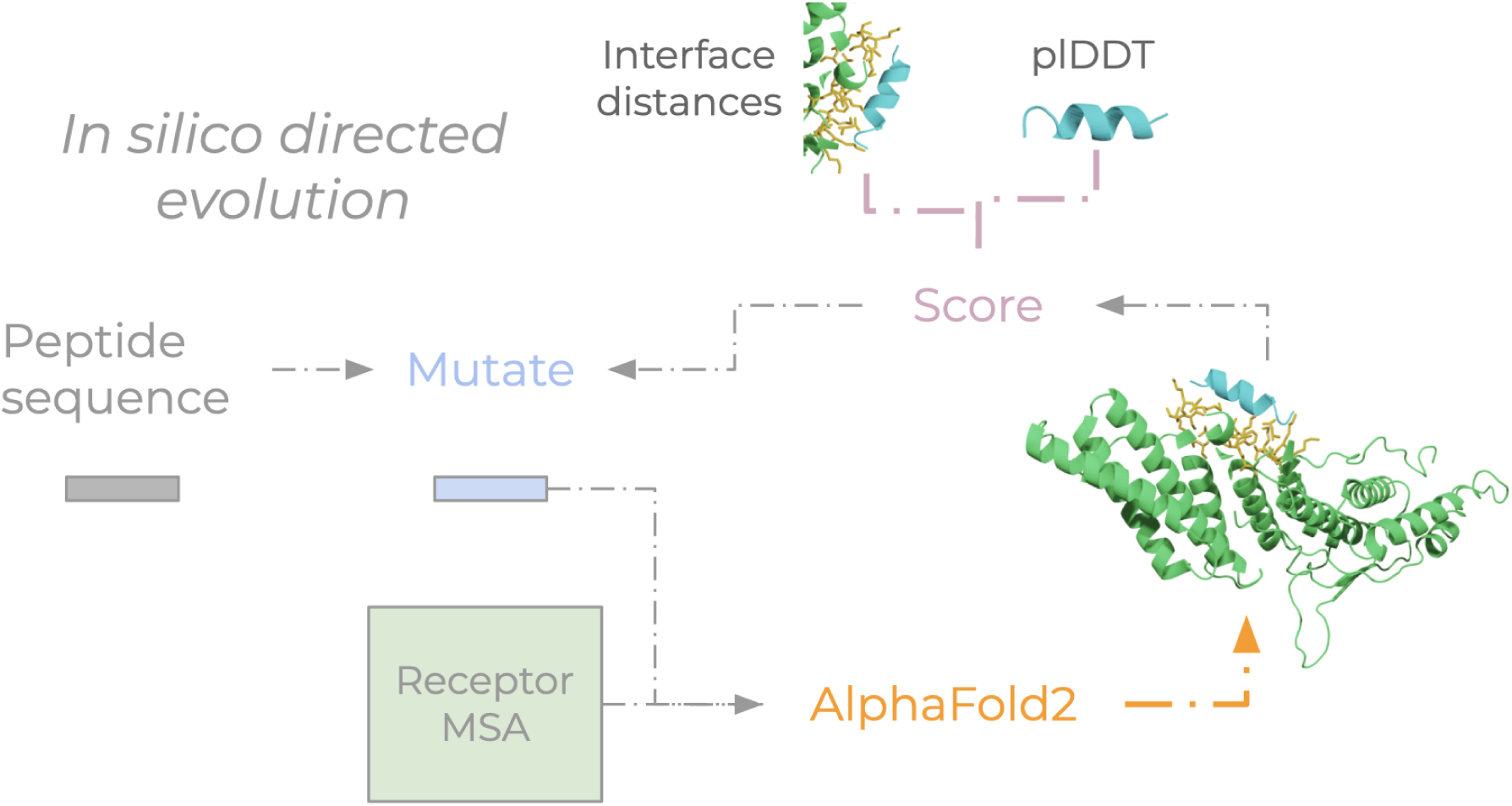
*In silico* directed evolution procedure towards an unspecified binding site. By not specifying a binding site, EvoBind2 is free to generate a structure-sequence combination to any part of the target protein as long as 1. the confidence (plDDT) of the predicted peptide is high and 2. the predicted peptide atom positions are close to the receptor surface (interface distances).

Where the peptide plDDT is the average plDDT over the peptide and dj is the shortest distance between all atoms n in the peptide and any atom in the target receptor.

The receptor is represented by a multiple sequence alignment that is constructed from searching uniclust30_2018_08[30] with HHblits[31] version 3.1.0:

`hhblits -E 0.001 -all -oa3m -n 2`

The peptide is represented by a single sequence (initialised with a Gumbel distribution over all standard 20 amino acids). The single chain AlphaFold2 folding pipeline [1] with model_1, one ensemble and 8 recycles was used. The optimisation is run for 1000 iterations and repeated five times from random starting points (sequences).

### Other binder sequence design methods

#### MC and MCTS with AlphaFold2 modifications

There are other methods[16] for designing binders using the loss function from the previously developed EvoBind [15], but no binder design method is entirely untargeted (i.e. a target site is required) and purely sequence-based (i.e. a target scaffold is needed).

#### Design with inverse folding

Inverse folding has been applied to the problem of binder design [14,32]. However, these methods require scaffolds that are not available for novel targets and have only been applied to design larger protein structures (not peptides) [6,7,33]. Other methods have recently been developed to predict and design peptide structures [34], but not in complex with a target protein and how to generate sequences for these is not clear. In addition, sequences for only 4-7% of structural seeds can be redesigned into sequences that structure prediction networks understand using inverse folding [10]. Relaxing the entire sequence-structure space will allow for more possibilities, increasing the probability that a structure prediction network can understand a designed sequence-structure combination.

### *In silico* evaluation with AlphaFold-multimer to avoid adversarial designs

In the EvoBind2 protocol developed here, no target site is used as this information is not available for novel target proteins. To ensure the design outcomes from the untargeted protocol are viable we use AlphaFold-multimer (AFM, **Figure 6**) as an additional check to avoid adversarial designs. This allows us to utilise the original EvoBind1 loss function (equation 2) which relies on a specified target site and a known peptide binder. In this case, we compare the loss between the two predictions using the EvoBind2 designs as “pseudo-native” structures.

**Figure 6.**
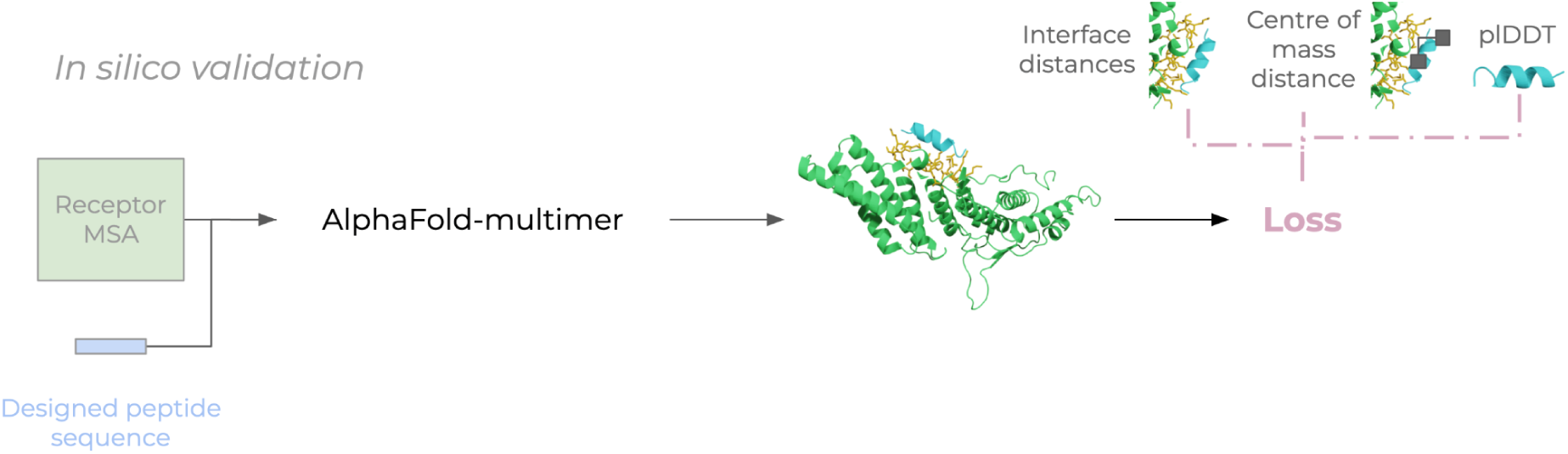
AFM validation. Given a receptor MSA and a single sequence for the designed peptide, AlphaFold-multimer is used to predict a protein-peptide complex. Based on the similarity of the predicted structure to that predicted with EvoBind2 and the confidence (plDDT) of the prediction a loss is calculated (equation 2).

Previous benchmarking has found that the best option for protein-peptide complex prediction is AFM v2.1.0 (params_model_1_multimer_v2) with 3 recycles and dropout everywhere but in the structural model (highest correlation with the ranking confidence from AFM+median DockQ) [12]. To account for that some examples improve with more recycles, we set the number of recycles to 20 and include ‘early stopping’ (no more recycles are included if the confidence score is not improved).

Calculating the loss using EvoBind2 vs. AFM predictions circumvents the need to know the native structure or CA centre of mass to use the EvoBind loss function. The EvoBind loss function, which has demonstrated accuracy both *in silico* and *in vitro* [15,16], can thus be used to evaluate the agreement between different neural networks (AF2 and AFM).

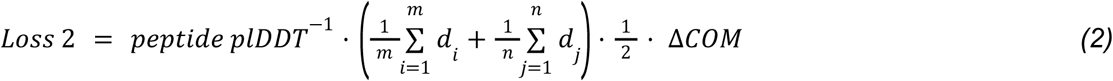

Where the peptide plDDT is the average plDDT from AFM over the peptide, di is the shortest distance between all atoms in the receptor target atoms (Cβs within 8 Å between the peptide and the receptor protein as predicted by EvoBind2) m and any atom in the peptide, dj is the shortest distance between all atoms n in the peptide and any atom in the receptor target atoms and ΔCOM is the CA centre of mass distance between the predicted peptides from the design and validation procedures, respectively.

### Binder design selection

We select the designs with the best combined loss calculated as the sum of equations 1 and 2. For the adversarial selection, we select designs with an AFM loss >1 according to equation 2 and the lowest possible loss according to equation 1, obtaining losses in the range 0.04-0.1 with equation 1 for the linear design task.

### Cyclic offset

To design cyclic binders, we implement a cyclic offset [19] informing the structure prediction network to connect the peptide amino acids in a continuous cycle. This feature is called the relative positional encoding in the AlphaFold2 [1] and AlphaFold-multimer [2] networks.

### PAE

The predicted aligned error (PAE) is a predicted confidence metric from the AlphaFold-mulitmer network that provides an estimate of the 2D distance error between residues [2]. Here, we extracted the PAE in the protein-peptide interface and averaged it to compare its relationship with the affinity (Kd) measured with SPR (Supplementary Figure 7).

### Positive control

To ensure that the expressed protein (semi-synthetic Ribonuclease, PDB ID 1SSC: https://www.rcsb.org/structure/1ssc) is in a structural state where it can interact with designed binders we analysed the affinity of the known peptide binder (11 residues) with sequence: PYVPVHFDASV. Using single-cycle kinetics with surface plasmon resonance (SPR) we obtained a Kd of 35 nM (Supplementary Figure 6). At the same time, many of the adversarial designs report no response suggesting that the protein is in a selective structural state where only specific peptide sequences will bind with high affinity.

### Length and initialisation

Figure 7 shows the best EvoBind2 loss (equation 1) vs the binder length for linear and cyclic design across five different runs (initialisations). For the linear design, the loss increases with the length, but no such relationship is apparent for the cyclic design. The losses vary substantially between runs (Supplementary Figures 1 and 2), but the results show that given five tries comparable results can be obtained in terms of affinity for linear designs. The finding that longer lengths display higher best losses and larger variability may be related to the higher number of possibilities accompanied by a longer sequence length making it less likely to end up in a favourable design.

**Figure 7.**
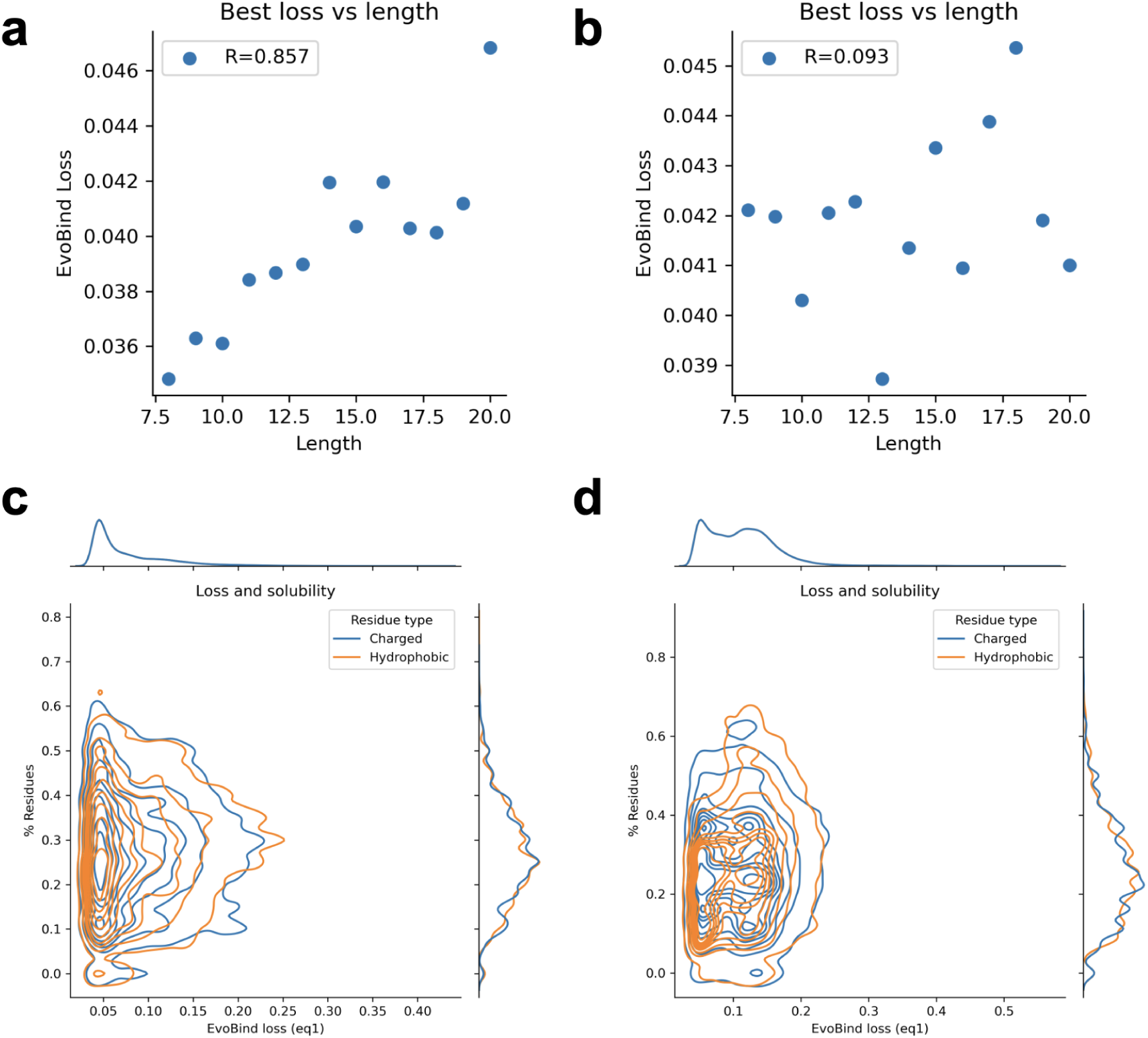
Best EvoBind2 loss (equation 1) vs length for linear (a) and cyclic (b) design. For the linear design, the loss increases with the length (Spearman R=0.857), but for the cyclic, such a relationship is not as apparent (Spearman R=0.093). c, d Fraction of charged (D, K, R, H and E) and hydrophobic residues (W, L, I, F, M, V, Y) vs EvoBind2 loss (equation 1) for linear (a) and cyclic (b) design. Different fractions of charged/hydrophobic residues are observed for a variety of losses suggesting that the solubility is not important for EvoBind2 when generating designs.

### Solubility

To ensure water solubility we select designed binders that have an average plDDT>90, >25% charged residues (D, K, R, H and E) and <25% hydrophobic residues (W, L, I, F, M, V, Y). Figures 6c and d show different fractions of charged/hydrophobic residues are observed for a variety of losses suggesting that solubility is not important for EvoBind2 when generating designs. All synthesised peptides were soluble in the SPR running buffer (see Affinity measurement with SPR).

### Protein expression and purification

The psfRNAseA-c001 construct was transformed into E. coli BL21 (DE3) T1R cells and cultivated in Terrific Broth (TB) medium supplemented with glycerol, ampicillin, and chloramphenicol. The cultures were grown at 37°C, and protein expression was induced with IPTG at an OD of approximately 3, after which the temperature was reduced to 18°C for overnight expression. The cells were harvested, resuspended in an IMAC lysis buffer, and disrupted by sonication. The soluble fractions were purified using IMAC and size exclusion chromatography, and the target protein was pooled, concentrated, and stored at −80°C. The psfRNAseA protein eluted together with some contaminants or possibly proteolytic fragments, and the full-length psfRNAseA protein with the MBP-tag intact was identified by SDS-PAGE analysis.

### Peptide synthesis

Lyophilized powders of peptides designed by EvoBind2 were synthesised and purified by JPT Peptide Technologies GmbH (linear peptides) and GenScript (cyclic peptides). For the top-selected cyclic peptide of length 17 (Supplementary Table 3), GenScript attempted several rounds of synthesis but were unable to produce the peptide. The quality of successfully synthesised peptides (purity above 90%) was verified by high-performance liquid chromatography and mass spectrometry. Peptides were resuspended in HBS-P+ Buffer (0.01 M HEPES, 0.15 M NaCl, and 0.05% v/v Surfactant P20) to achieve a stock concentration of 10 mM.

### Affinity measurement with SPR

To directly measure the binding affinity of designed peptides toward 1SSC we used surface plasmon resonance (SPR). An experimental method that measures real-time binding interactions between biomolecules by detecting changes in refractive index near a sensor surface. All interaction analysis was performed at 25 °C in a running buffer of HBS-P+ (10 mM HEPES, pH 7.4, 150 mM NaCl, 0.05% Tween-20) for Biacore 8K. MBP-8xHis-1SCC was immobilised to about 12000 RU using standard amine coupling reagents 1-ethyl-3-(3-dimethylaminopropyl)carbodiimide (EDC) and N-hydroxysuccinimide (NHS) onto a Series S Sensor Chip CM5. The designed peptides were captured onto the prepared surface to generate an interaction with a maximum response (R_max_) from 20 RU to 300 RU.

Two independent single-cycle kinetics experiments were conducted using different concentration ranges of the peptides as the analytes. In the first experiment, three concentrations of the analyte were used: 2 μM, 10 μM, and 50 μM. The second experiment employed a five-point concentration series: 3.125 μM, 6.25 μM, 12.5 μM, 25 μM, and 50 μM. This dual-range approach was chosen to provide a broader dynamic range for capturing both low and high-affinity interactions, while also allowing for more precise Kd determination within a narrower, finely graduated range. All dilutions were prepared in the running buffer.

Each concentration series was injected in a single cycle without regeneration steps between injections. The flow rate was set at 30 μL/min, with an association time of 120 seconds and a dissociation time of 1800 seconds. Data processing and analysis were performed using Biacore Insight Evaluation Software. Raw data were preprocessed by subtracting reference data from sample data. Sensorgrams were then fitted to a 1:1 binding model (equations 3 and 4) [35] using global kinetic rate constants for association (ka) and dissociation (kd), along with R_max_ values per sample. This selection process was applied to both experiments, resulting in a final dataset comprising high-quality fits from both the three-concentration and five-concentration series. The retained data were used to determine the association rate constant (ka), dissociation rate constant (kd), and equilibrium dissociation constant (KD). See Supplementary Figures 4-7 for all sensorgrams.

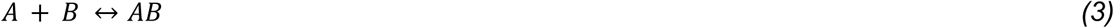

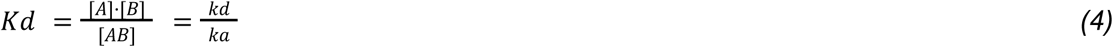

Where A is the protein receptor that was immobilised on the sensor chip, and B is the analyte (peptide). The equilibrium dissociation constant Kd is calculated as the ratio of kd to ka, where ka represents the association rate constant for complex formation (A+B →AB, M^-1^s^-1^), and kd the dissociation rate constant (AB →A+B, s^-1^).

## Supplementary information

### Supplementary data

**Supplementary Table 1.**
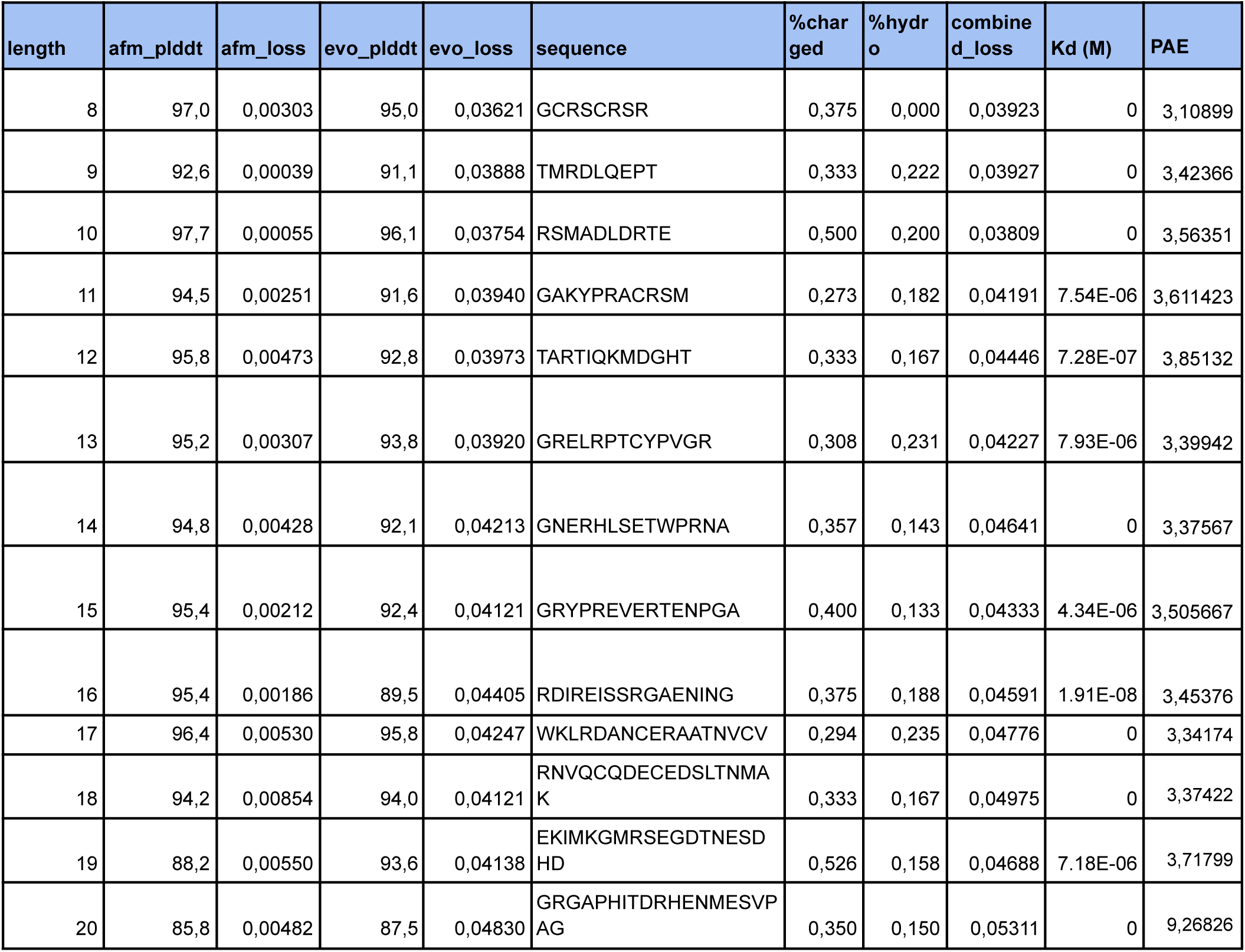
Linear binder design selection. One sequence was selected per length according to the lowest combined loss. The ‘length’ column represents the length of the selected peptide sequences. ‘afm_plddt’ and ‘evo_plddt’ are the AlphaFold-multimer and EvoBind2 pLDDT (per-residue model confidence) scores respectively, which are measures of the predicted accuracy of the protein structure. ‘afm_loss’ (equation 2) and ‘evo_loss’ (equation 1) are the loss values associated with the AFM and EvoBind2 predictions respectively. The ‘sequence’ column contains the amino acid sequence of the protein. ‘%charged’ and ‘%hydro’ show the percentage of charged and hydrophobic amino acids in the sequence respectively. The ‘combined_loss’ (sum of equations 1 and 2) is a combined measure of loss, considering both the AFM and EvoBind2 predictions. The kinetic properties of all the sequences towards the receptor protein were measured in the wet lab using Biacore 8K SPR and data are listed in the ‘Kd’ column.

**Supplementary Table 2.**
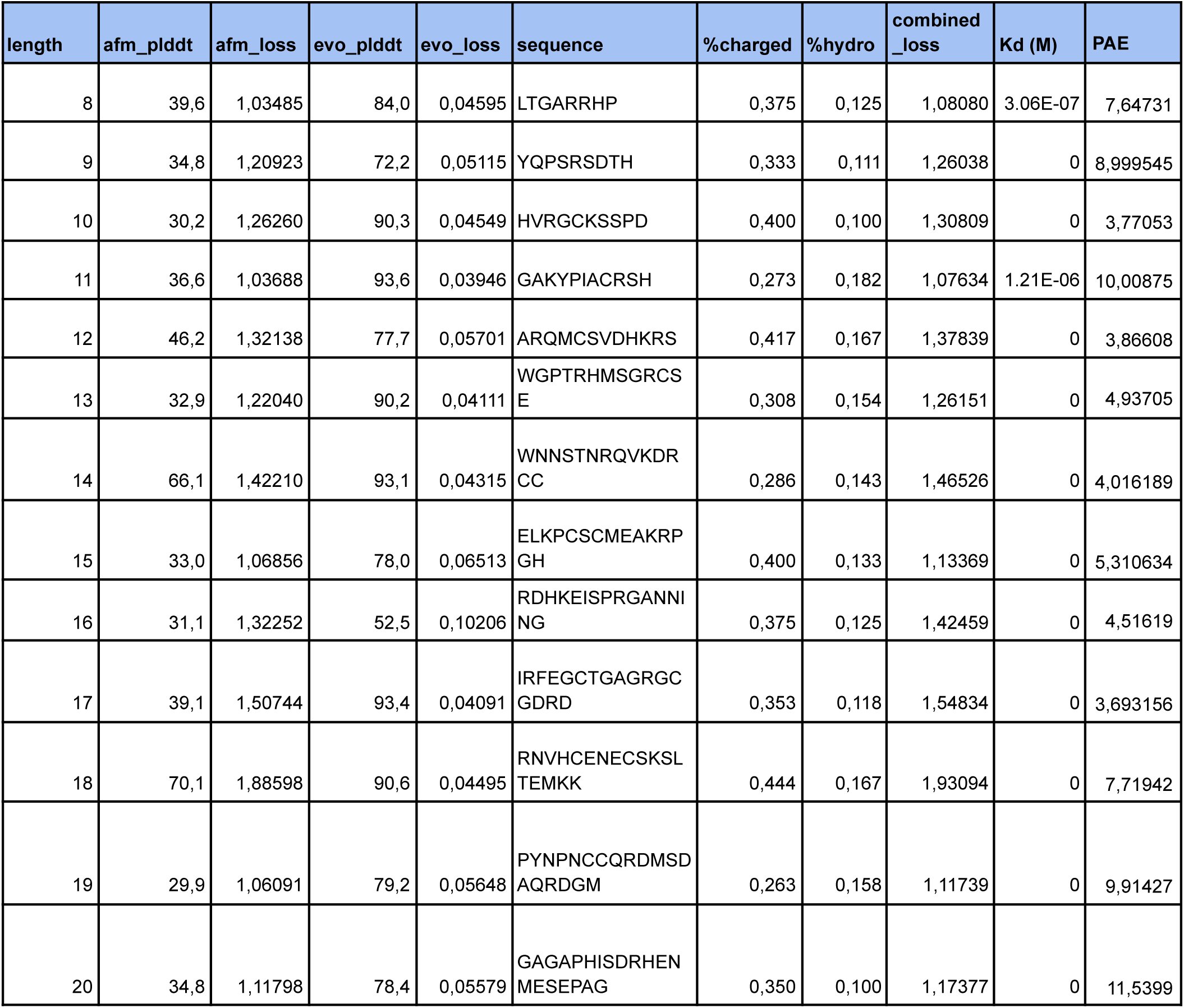
Adversarial linear design selection. One sequence was selected per length according to having an AFM loss >1 and lowest evobind loss. The ‘length’ column represents the length of the adversarial linear peptide sequences. ‘afm_plddt’ and ‘evo_plddt’ are the AlphaFold-multimer and EvoBind2 pLDDT (per-residue model confidence) scores respectively, which are measures of the predicted accuracy of the protein structure. ‘afm_loss’ (equation 2) and ‘evo_loss’ (equation 1) are the loss values associated with the AFM and EvoBind2 predictions respectively. The ‘sequence’ column contains the amino acid sequence of the protein. ‘%charged’ and ‘%hydro’ show the percentage of charged and hydrophobic amino acids in the sequence respectively. The ‘combined_loss’ (sum of equations 1 and 2) is a combined measure of loss, considering both the AFM and EvoBind2 predictions. The kinetic properties of all the sequences towards the receptor protein were measured in the wet lab using Biacore 8K SPR and data are listed in the ‘Kd’ column.

| length | afm_plddt | afm_loss | evo_plddt | evo_loss | sequence | %charged | %hydro | combined_loss | Kd (M) | PAE |
| --- | --- | --- | --- | --- | --- | --- | --- | --- | --- | --- |
| 8 | 39,6 | 1,03485 | 84,0 | 0,04595 | LTGARRHP | 0,375 | 0,125 | 1,08080 | 3.06E-07 | 7,64731 |
| 9 | 34,8 | 1,20923 | 72,2 | 0,05115 | YQPSRSDTH | 0,333 | 0,111 | 1,26038 | 0 | 8,999545 |
| 10 | 30,2 | 1,26260 | 90,3 | 0,04549 | HVRGCKSSPD | 0,400 | 0,100 | 1,30809 | 0 | 3,77053 |
| 11 | 36,6 | 1,03688 | 93,6 | 0,03946 | GAKYPIACRSH | 0,273 | 0,182 | 1,07634 | 1.21E-06 | 10,00875 |
| 12 | 46,2 | 1,32138 | 77,7 | 0,05701 | ARQMCSVDHKRS | 0,417 | 0,167 | 1,37839 | 0 | 3,86608 |
| 13 | 32,9 | 1,22040 | 90,2 | 0,04111 | WGPTRHMSGRCSE | 0,308 | 0,154 | 1,26151 | 0 | 4,93705 |
| 14 | 66,1 | 1,42210 | 93,1 | 0,04315 | WNNSTNRQVKDRCC | 0,286 | 0,143 | 1,46526 | 0 | 4,016189 |
| 15 | 33,0 | 1,06856 | 78,0 | 0,06513 | ELKPCSCMEAKRPGH | 0,400 | 0,133 | 1,13369 | 0 | 5,310634 |
| 16 | 31,1 | 1,32252 | 52,5 | 0,10206 | RDHKEISPRGANNI NG | 0,375 | 0,125 | 1,42459 | 0 | 4,51619 |
| 17 | 39,1 | 1,50744 | 93,4 | 0,04091 | IRFEGCTGAGRGC GDRD | 0,353 | 0,118 | 1,54834 | 0 | 3,693156 |
| 18 | 70,1 | 1,88598 | 90,6 | 0,04495 | RNVHCENECSKSL TEMKK | 0,444 | 0,167 | 1,93094 | 0 | 7,71942 |
| 19 | 29,9 | 1,06091 | 79,2 | 0,05648 | PYNPNCCQRDMSD AQRDGM | 0,263 | 0,158 | 1,11739 | 0 | 9,91427 |
| 20 | 34,8 | 1,11798 | 78,4 | 0,05579 | GAGAPHISDRHEN MESEPAG | 0,350 | 0,100 | 1,17377 | 0 | 11,5399 |

**Supplementary Table 3.**
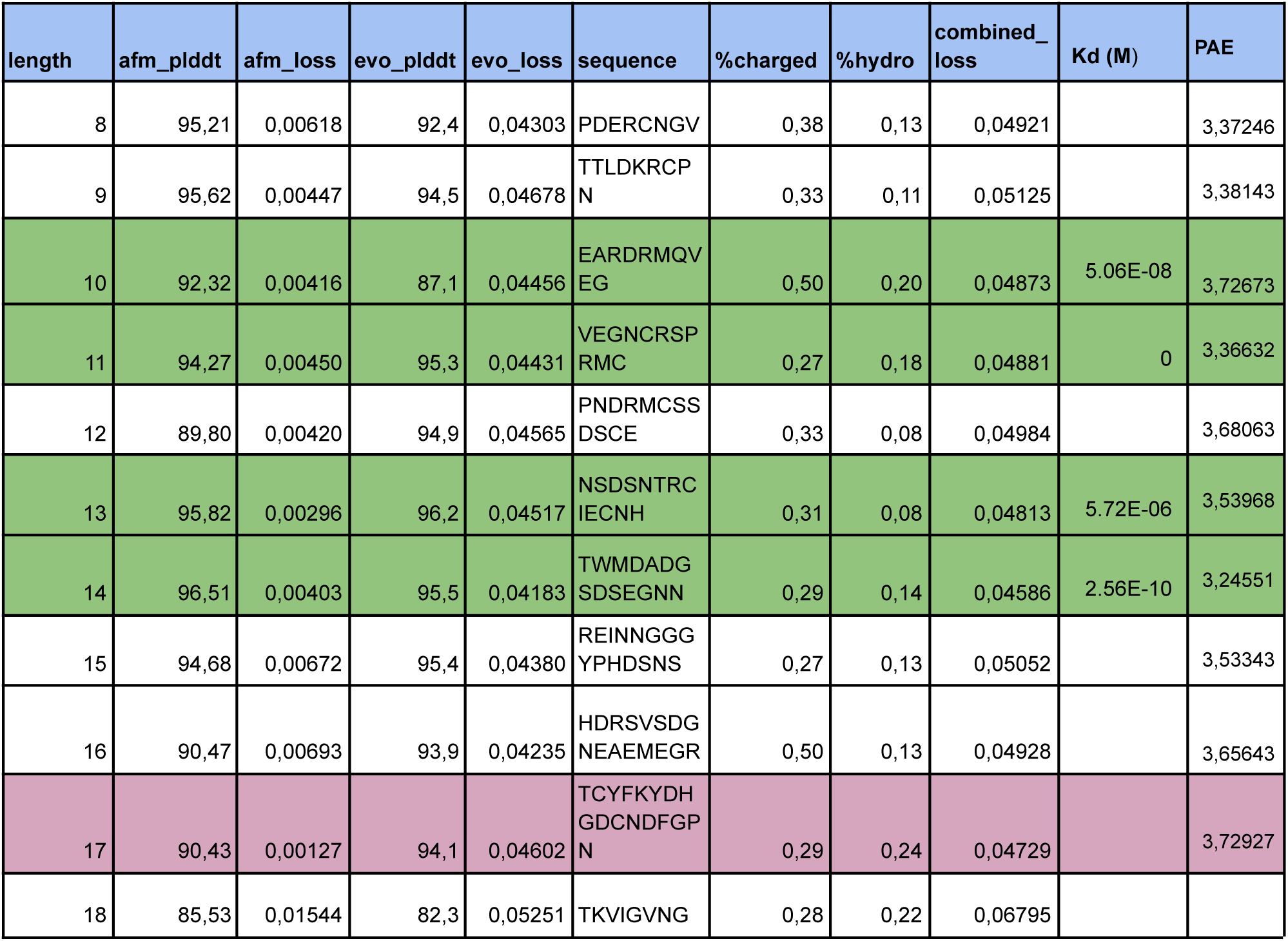
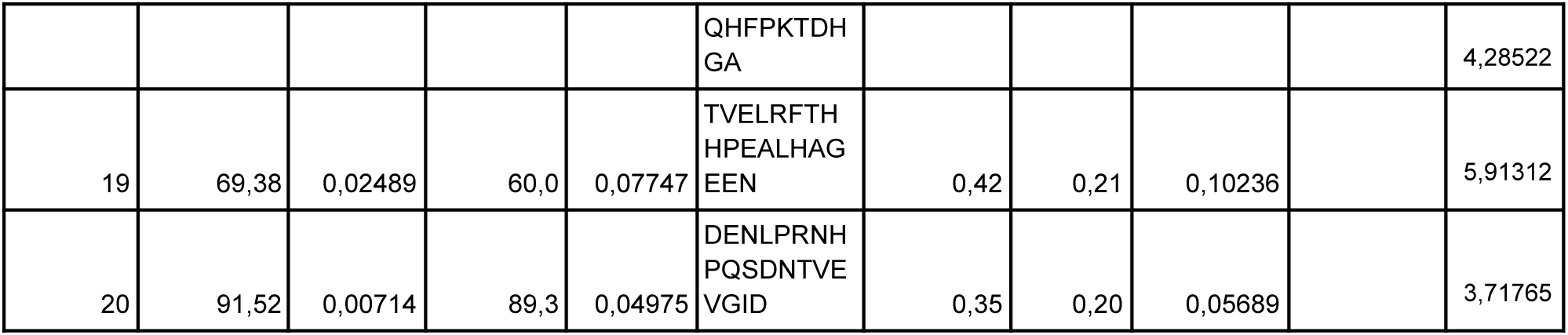
Cyclic binder design selection. One sequence was selected per length according to the lowest combined loss. The ‘length’ column represents the length of the cyclic peptide sequences. ‘afm_plddt’ and ‘evo_plddt’ are the AlphaFold-multimer and EvoBind2 pLDDT (per-residue model confidence) scores respectively, which are measures of the predicted accuracy of the protein structure. ‘afm_loss’ (equation 2) and ‘evo_loss’ (equation 1) are the loss values associated with the AFM and EvoBind2 predictions respectively. The ‘sequence’ column contains the amino acid sequence of the protein. ‘%charged’ and ‘%hydro’ show the percentage of charged and hydrophobic amino acids in the sequence respectively. The ‘combined_loss’ (sum of equations 1 and 2) is a combined measure of loss, considering both the AFM and EvoBind2 predictions. The lengths selected for experimental validation (top 4) are coloured green for the ones that were evaluated. Length 17 could not be synthesised by GenScript and is therefore coloured magenta. The remaining peptides were not evaluated experimentally.

### Supplementary figures

**Supplementary Figure 1.**
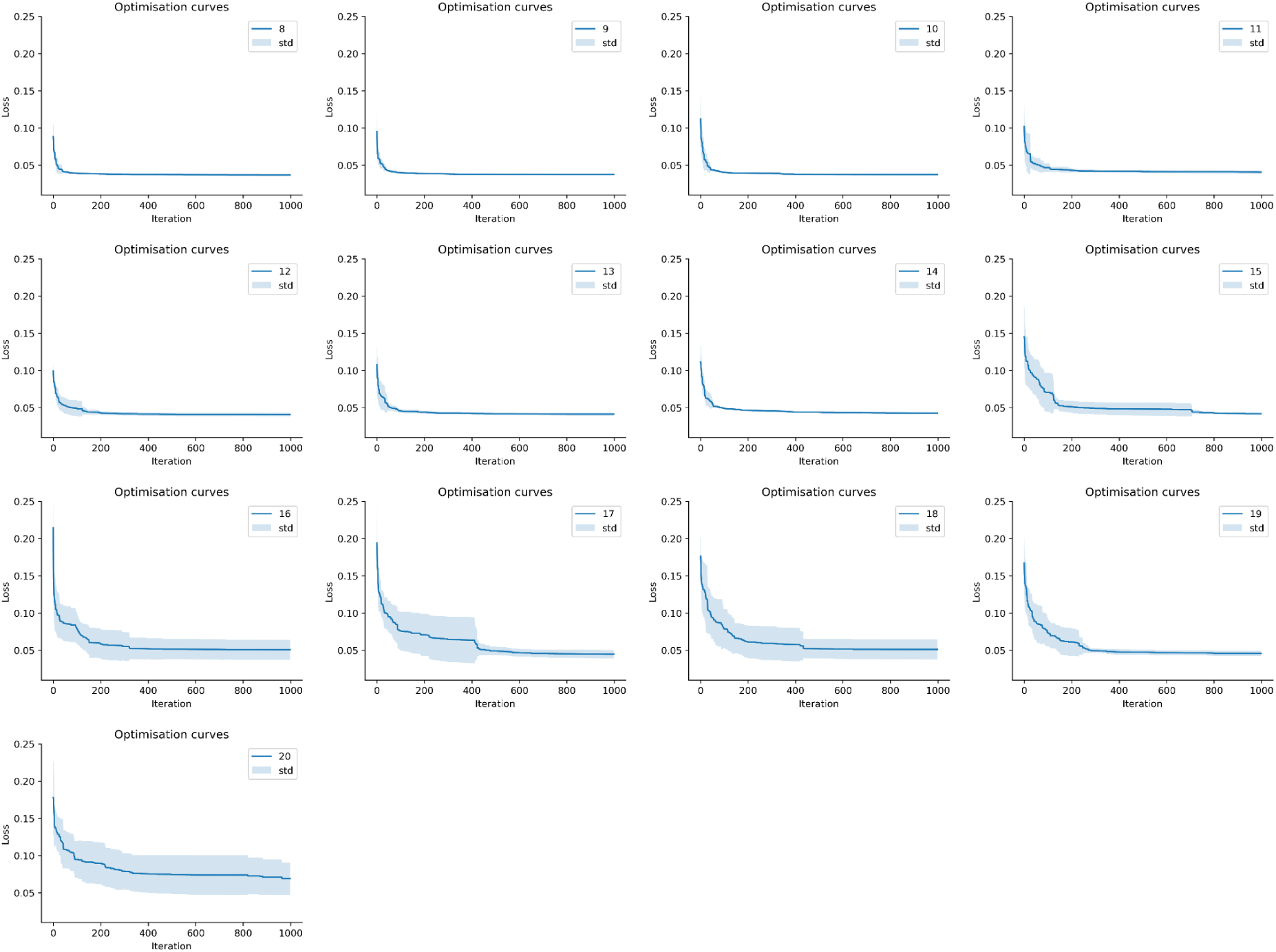
Linear optimisation. EvoBind2 loss (equation 1) vs. Iteration curve showing the optimisation process 1000 iterations. The blue line represents the mean loss value for each peptide length across five different initialisations, and the shaded region is the standard deviation. The standard deviation increases with the length, suggesting the higher number of possibilities accompanied by a longer sequence length making it less likely to end up in a favourable design.

**Supplementary Figure 2.**
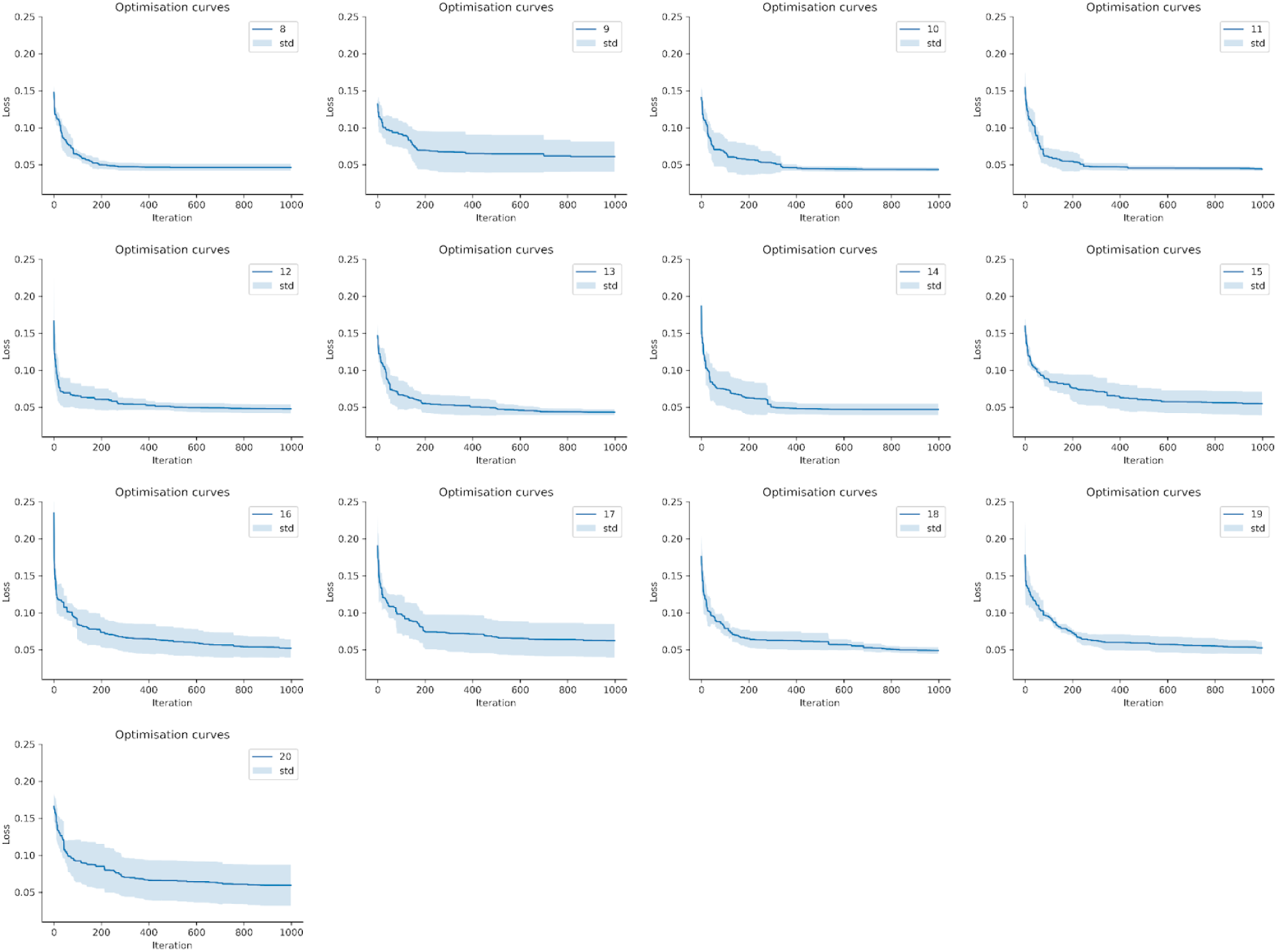
Cyclic optimisation. EvoBind2 loss (equation 1) vs. Iteration curve showing the optimisation process 1000 iterations. The blue line represents the mean loss value for each peptide length across five different initialisations, and the shaded region is the standard deviation. The standard deviation does not increase with the length as with the linear selection.

**Supplementary Figure 3.**
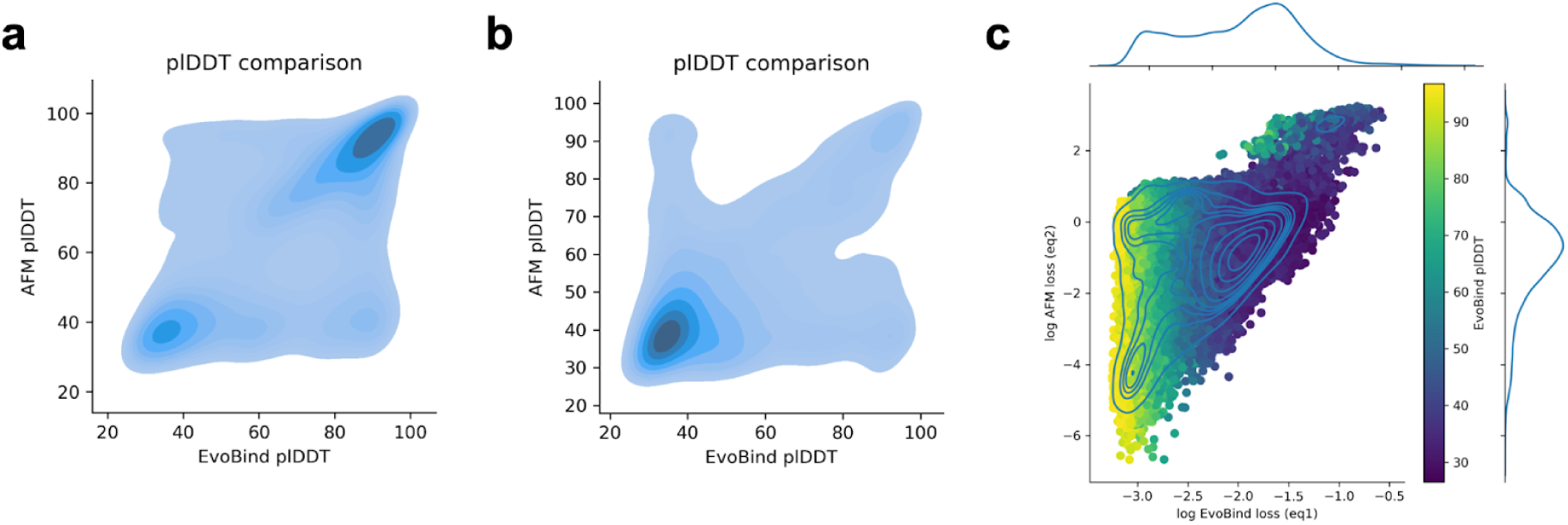
plDDT comparison for linear (a) and cyclic design (b). c) Adversarial comparison for cyclic peptide design. EvoBind2 loss (equation 1) vs AlphaFold-multimer (AFM) loss (equation 2) on log scale for all cyclic design runs (n=64935). The points are coloured by the plDDT from EvoBind2 and the density is represented with blue lines.

**Supplementary Figure 4.**
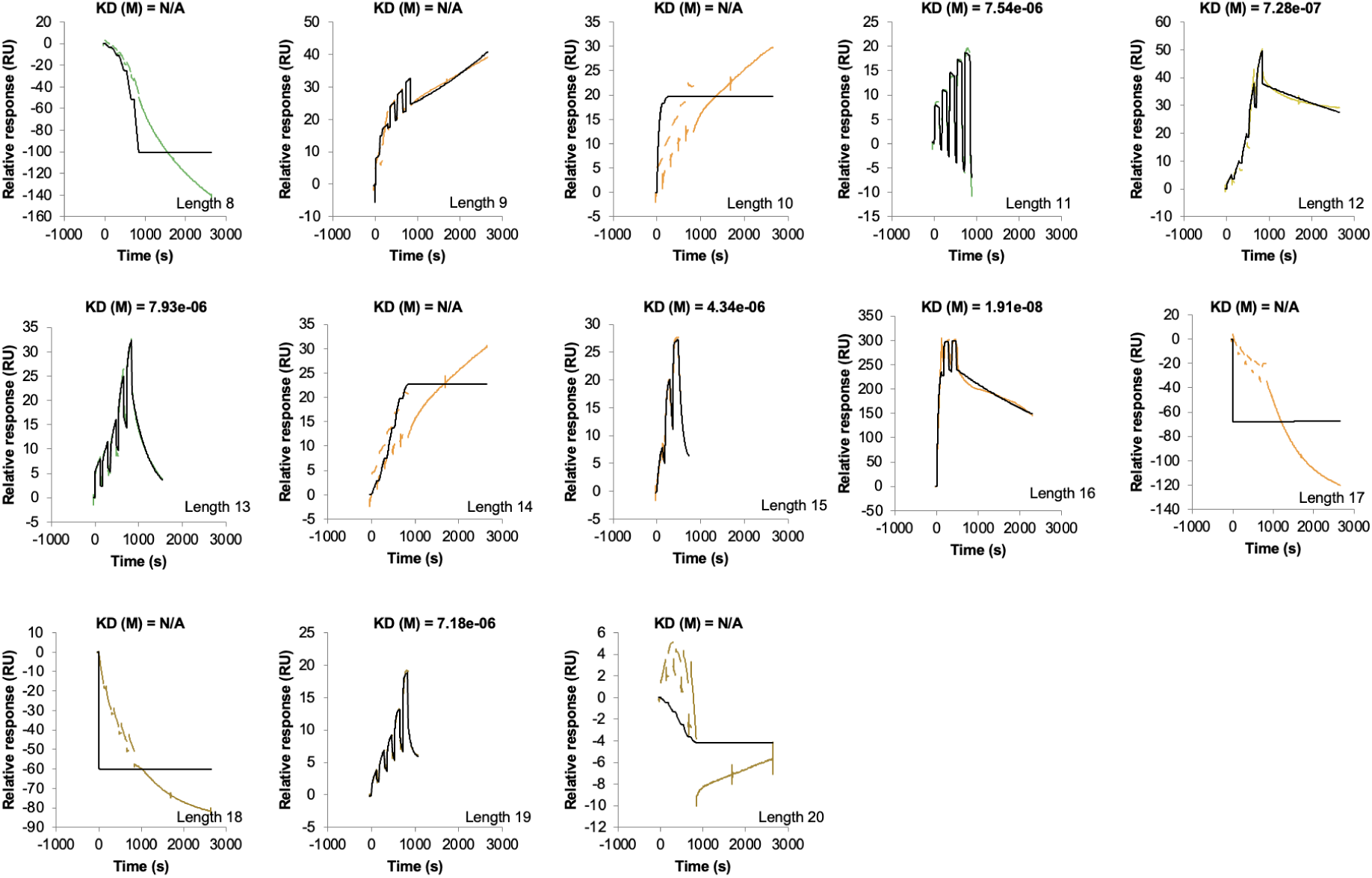
Single cycle kinetic SPR curves for the top linear selection. Relative Response (RU) vs Time (s) for designed peptides of different lengths, showing the association and dissociation phases of the binding interaction. Kd values are shown where measurable; ‘N/A’ denotes affinity beyond the assay’s detection limit. Raw data appears in black, while green lines represent the curve fit used for the kinetic analysis. The highest affinity is obtained for length 16 (19 nM) and the lowest for length 13 (7.93 μM). In total, 6/13 binders display affinity resulting in a 46% success rate.

**Supplementary Figure 5.**
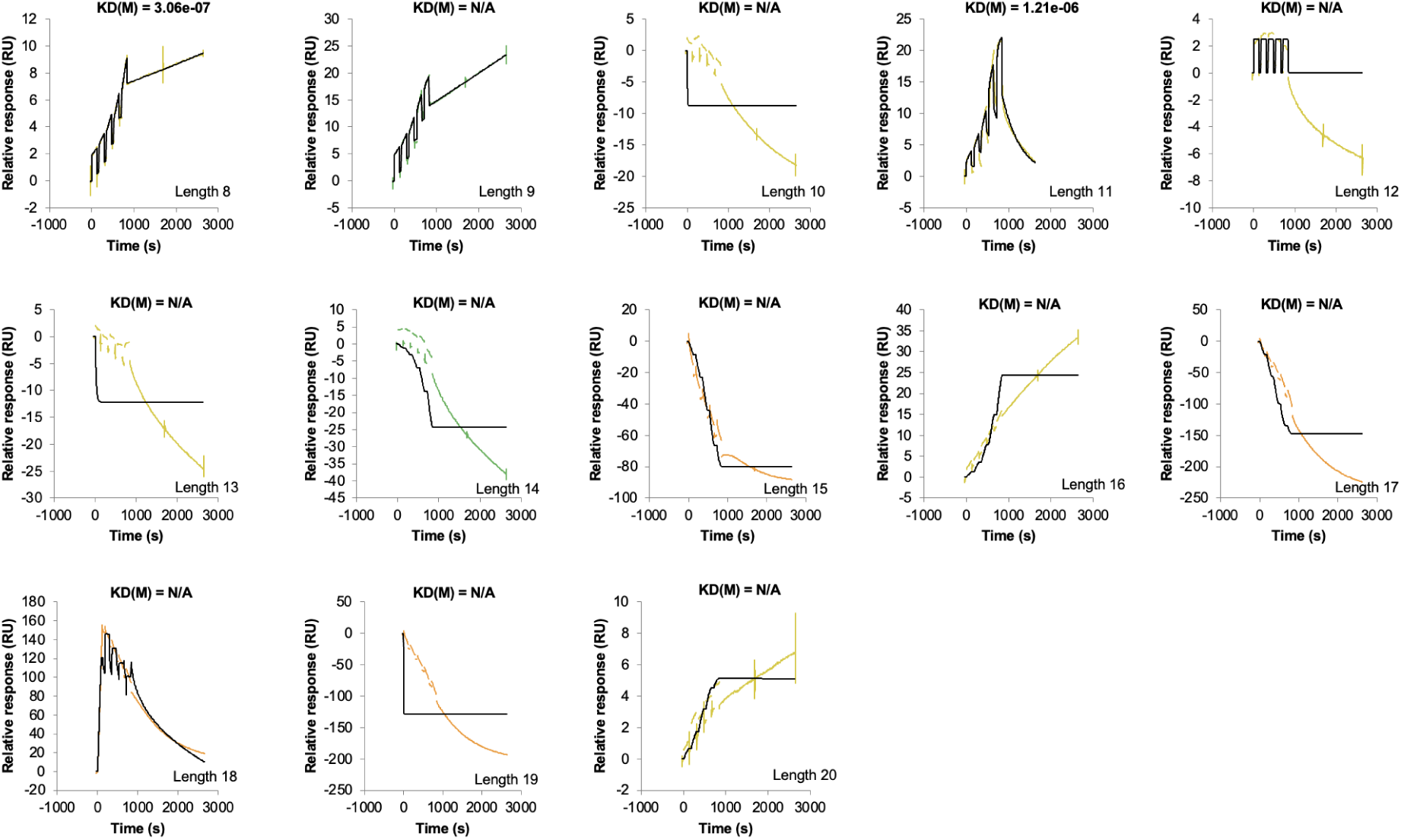
Single cycle kinetic SPR curves for the adversarial selection. Relative Response (RU) vs Time (s) for designed peptides of different lengths, showing the association and dissociation phases of the binding interaction. Kd values are shown where measurable; ‘N/A’ denotes affinity beyond the assay’s detection limit. Raw data appears in black, while green lines represent the curve fit used for the kinetic analysis. Compared to the top selection (Supplementary Figure 4), only 2 peptides display measurable affinity (lengths 8 and 11, 15% success rate).

**Supplementary Figure 6.**
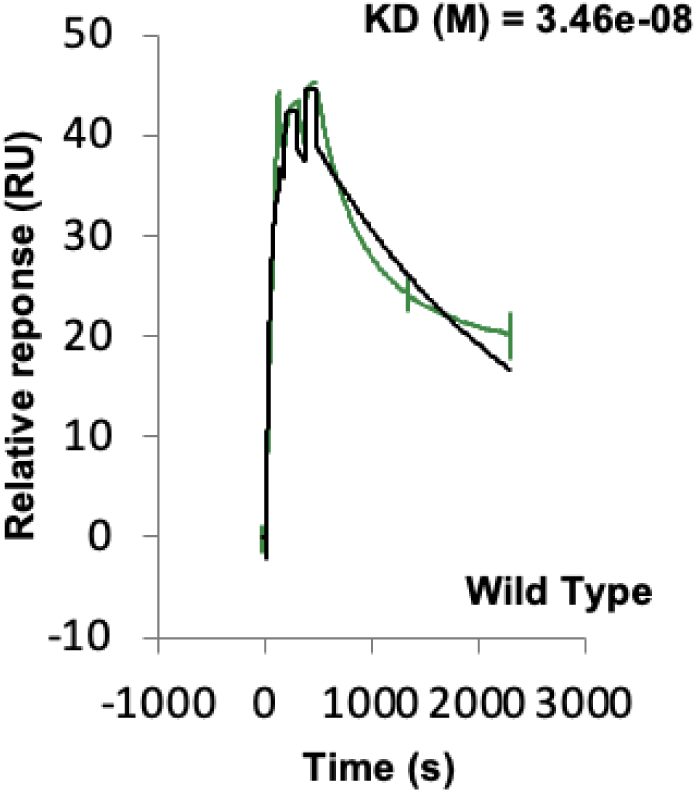
Single cycle kinetic SPR curve (Relative Response (RU) vs Time (s)) for the “wild-type” peptide used for the semi-synthetic Ribonuclease, PDB ID 1SSC: https://www.rcsb.org/structure/1ssc). The affinity (KD) is 25 nM, suggesting expected behaviour from this positive control and that the target protein is in a functional state.

**Supplementary Figure 7.**
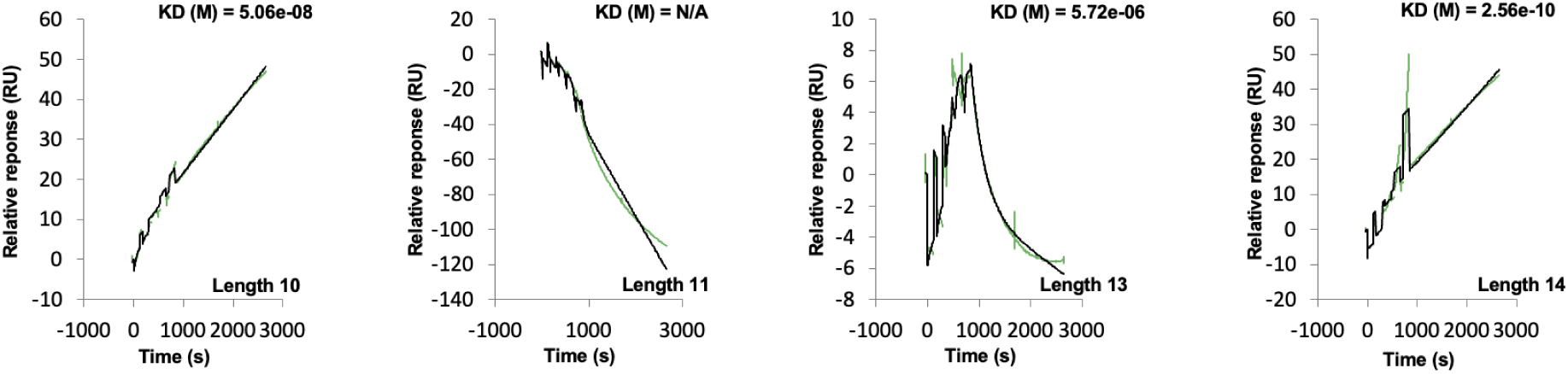
Single cycle kinetic SPR curves for the top cyclic selection. Relative Response (RU) vs Time (s) for four selected designed cyclic peptides of different lengths, showing the association and dissociation phases of the binding interaction. Kd values are shown where measurable; ‘N/A’ denotes affinity beyond the assay’s detection limit. Raw data appears in black, while green lines represent the curve fit used for the kinetic analysis. Compared to the linear selection (Supplementary Figure 4) and the WT (Supplementary Figure 6), the highest affinity (0.26 nM, length 14) is over 100x stronger suggesting that cyclic peptides can be made to outperform linear ones. In addition, 3 out of 4 binders display affinity (75 % the success rate) compared to 46% for the linear designs.

**Supplementary Figure 8.**
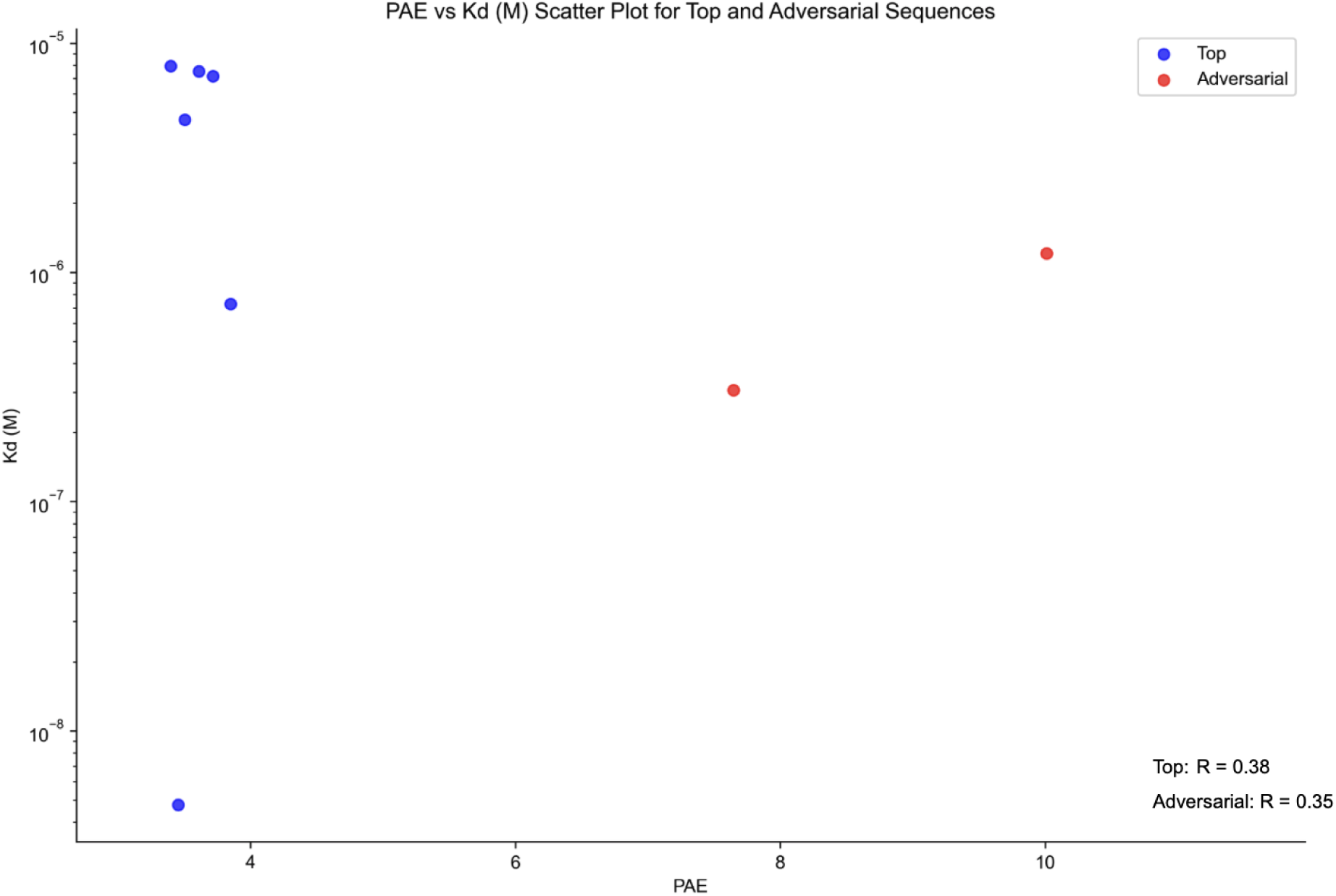
This scatter plot compares the Predicted Aligned Error (PAE) against the dissociation constant (Kd) for designed linear top selected and adversarial peptide binders. Blue points represent top designs, which cluster at lower PAE values (3-4) and red points show adversarial designs with higher PAE (8-10), consistent with the AFM losses (Figure 3b). The Spearman correlations (R) are low suggesting no meaningful relationship between PAE and Kd.

## Notes

https://zenodo.org/records/13913345

### Competing Interest Statement

The authors have declared no competing interest.

### Summary of Updates

We have added the experimental results for the cyclic binders (they show sub nM affinity).

